# PlexinD1 deficiency in CD11c+ dendritic cells exacerbates airway hyperresponsiveness and enhances IgE and mucus production in allergic asthma

**DOI:** 10.1101/2023.09.11.557276

**Authors:** Lianyu Shan, Mojdeh Matloubi, Ifeoma Okwor, Sam Kung, Mohamed Sadek Almiski, Sujata Basu, Andrew Halayko, Latifa Koussih, Abdelilah S. Gounni

**Author notes:** Corresponding Author^a^: Abdelilah S. Gounni, Ph.D., Department of Immunology, 471 Apotex Centre 750 McDermot Avenue, University of Manitoba, Winnipeg, MB R3E 0T5 Canada. contributed equally to this work (Lianyu Shan, Mojdeh Matloubi). **Authorship:** Conception and design: A.S.G., L.S., M.M. Acquisition of data and analysis: L.S., M.M., I.O., S.B., L.K. Interpretation of data: L.S., M.M., I.O., M.S.A., S.B., L.K. Draft the article: M.M. Critically review the article: L.S., M.M., I.O., S.P.K., M.S.A., S.B., A.H., L.K., A.S.G. Final approval: L.S., M.M., I.O., S.P.K., M.S.A., S.B., A.H., L.K., A.S.G.

## Abstract

Dendritic cells (DC) play a crucial role in regulating allergic asthma. We have demonstrated that the absence of semaphorin3E (Sema3E) exacerbates asthma features in acute and chronic asthma models. However, the role of plexinD1 in these events, especially in DC is unknown. Therefore, we investigated the role of plexinD1 in CD11c+ DC in the HDM model of asthma. CD11c+ DC-specific plexinD1 knockout mice and wild-type mice were subjected to HDM acute allergen protocol. Airway hyperresponsiveness (AHR) parameters were measured using the FlexiVent ventilator. Lung tissue and bronchoalveolar lavage fluid (BALF) were processed by flow cytometry. Cytokines and antibodies were measured using mesoscale and ELISA. Collagen deposition and mucus production were visualized by histological staining, and associated genes were investigated using Real-time PCR. We showed that DC-specific plexinD1 knockout mice exhibited exacerbated airway hyperresponsiveness, including increased airway resistance and tissue elastance. These mice displayed enhanced levels of mucus production and collagen gene expression compared to wild-type mice. These events were accompanied by enhanced recruitment of conventional DCs, specifically CD11b+ cDC2, into the lungs and higher levels of total and HDM-specific serum IgE in *CD11c^PLXND1^ ^KO^* compared to wild-type counterparts. Mechanistically, a significantly higher level of IgE in the co-culture of B-DCs isolated from *CD11c^PLXND1^ ^KO^* mice compared to DCs isolated from wild-type mice. Overall, our data reveals that the Sema3E-plexinD1 signalling pathway in CD11c+ DC is critical in modulating asthma features.

**Graphical Abstract:** 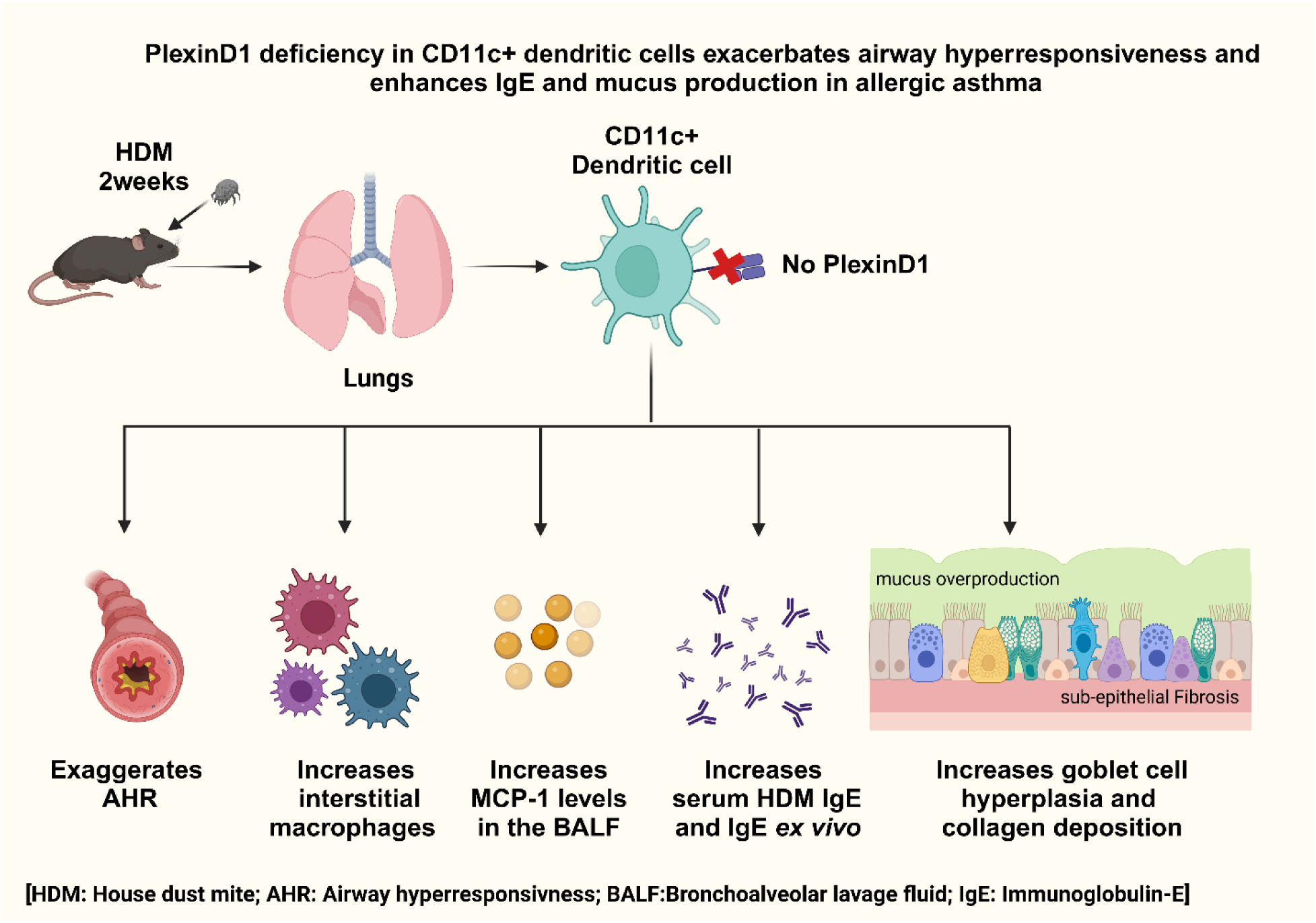

## Introduction

Asthma is a significant public health problem, affecting more than 358 million individuals worldwide (1). Allergic asthma is a chronic inflammatory disorder involving the airways, with a predominant Th2/Th17 immune response and an increase of innate and adaptive immune cells. Inflammatory cells release various mediators, such as cytokines, chemokines, histamine, immunoglobulins (Igs), growth factors, and lipid mediators, which contribute to the development of airway hyperreactivity, excessive mucus production, collagen deposition, hypertrophy and hyperplasia of airway smooth muscle (ASM), and subsequent changes in the airway architecture or remodelling (2). Despite advancements in drug therapies, approximately 5-10% of asthma patients remain unresponsive to these medications (3), highlighting the crucial need for novel therapeutic approaches and a deeper understanding of the underlying factors regulating the pathophysiology of asthma.

Semaphorins are discovered as an axon guidance cue during neural system development (4). However, they can be found in various organs and tissues and participate in different signaling pathways (5, 6). In pulmonary diseases, semaphorins play critical roles in cell-cell contact, cell migration, proliferation, differentiation, and immune system regulation (7–11). Among semaphorins, Semaphorin3E (Sema3E) has been investigated in asthma widely and appeared as a regulatory molecule (12–19). Sema3E binds to plexinD1 with high affinity as its canonical receptor (20–22). PlexinD1 has critical roles in axon guidance, vascular patterning, and regulation of double-positive thymocyte migration in the thymus (7, 22), and its loss of function is associated with autoimmune diseases and cancer (23).

Previous findings from our research group have indicated that Sema3E and its corresponding receptor, plexinD1, play critical roles in modulating airway inflammation, remodeling, and hyperresponsiveness during asthma (24). The global deletion of *sema3e* gene in mice led to high recruitment of granulocytes in the lungs and an increased Th2/Th17 immune response, accompanied by enhanced airway hyperresponsiveness (AHR), excessive mucus production, and collagen deposition or sub-epithelial fibrosis (24). Conversely, the administration of recombinant Sema3E intranasally ameliorated these pathological manifestations (24), thereby emphasizing the crucial homeostatic function of the Sema3E-plexinD1 axis within the context of allergic asthma.

Dendritic cells (DCs) connect innate and adaptive immune responses, acting as essential inflammatory cells (25). The functions of dendritic cells significantly influence immune system regulation in various inflammatory conditions. Therefore, investigating the mechanisms responsible for controlling dendritic cell functions impacts comprehending the initiation, progression, and disruption of immune responses (25).

Sema3E plays a significant role in modulating DC behavior in allergic asthma (17, 18). Using a mouse model of HDM-induced allergic asthma, we revealed that *Sema3E*-deficient (KO) mice exhibited an increased population of CD11c+ CD11b+ CD103-DCs, which consequently led to Th2/Th17 immune activation (18). Furthermore, HDM sensitization in Sema3E KO mice resulted in elevated expression of programmed death-ligand 2 (PDL2), interferon regulatory factor 4 (IRF-4) CC chemokine receptor-7 (CCR7) and enhanced allergen uptake capacity in pulmonary CD11b+ DCs, compared to their wild-type counterparts (17). *In vitro* experiments using Sema3E KO CD11c+ bone marrow-derived dendritic cells (BMDCs) demonstrated increased baseline migration associated with an enhanced Rac1 GTPase activity and actin polymerization in response to CCL21, in comparison to DCs isolated from wild-type mice. These findings suggest that the absence of Sema3E may augment the ability of DCs to uptake HDM allergens and migrate to the lymph nodes, subsequently leading to enhanced activation of T and B cells and further exacerbation of allergic asthma-associated inflammation (17). Mechanistically, our data revealed that in the absence of Sema3E, there was a substantial increase in IL-5, IL-13, IL-4, and IL-17A levels, along with a significant increase in both GATA3 and RORγt expression in CD4+ T cells from the mediastinal lymph nodes (mLN) of Sema3E KO mice following allergen challenge (18). These observations can account for the Th2/Th17-skewed phenotype observed in our HDM model of allergic asthma.

Moreover, upon the adoptive transfer of CD11b+ DCs from Sema3E-KO mice to wild-type mice, we observed the recapitulation of the abovementioned responses in the wild-type recipient mice (18). In light of these findings, we conclude that Sema3E acts as a guidance cue for recruiting lung DCs and modulates T and B cell responses in allergic asthma. Nevertheless, the role of plexinD1 in these events is not clear. Therefore, we investigated the role of plexinD1 deficient CD11c+ DC in a murine model of allergic asthma. In this study, we found that deficiency of plexinD1 in CD11c+ DC exacerbates airway hyperresponsiveness (AHR), enhances mucus production, and upregulates collagen gene expression. Moreover, the absence of plexinD1 in CD11c+ DC leads to the accumulation of conventional DC, particularly CD11b+ cDC2s, in the lungs, along with increased IgE levels and overall increased inflammation and tissue remodelling. These findings indicate that Sema3E-plexinD1 plays a crucial regulatory role in allergic asthma features.

## Method and materials

### Animals

*PlexinD1^fl/fl^* mice (B6;129-*Plxnd1^tm1.1Tmj/J^*) (10) were kindly provided by Dr. T.M. Jessell (Columbia University/Howard Hughes Medical Institute, New York, NY) and were crossed with *B6.Cg-Tg*(*Itgax-cre)^1-1Reiz/J^* mice (The Jackson Laboratory, stock number 008068). The latter mice express Cre recombinase under the control of the mouse integrin alpha X (CD11c) promoter, resulting in the generation of *CD11c(Itgax-cre):PlxnD1^fl/fl^* mouse. All mice were housed in the pathogen-free room at the Central Animal Care facility, University of Manitoba. All procedures followed the guidelines provided by the Canadian Council for Animal Care and approved by the University of Manitoba Animal Care and Use Committee (protocol number 19-035).

### HDM-induced airway inflammation model

Six- to eight-week-old female *PLXND1^fl/fl^* (wild-type) and *CD11c^PLXND1^ ^KO^* mice received intranasal administration of 25 mg of HDM extract (lot 259585; Greer Laboratories, Lenoir, NC) in 35μl of sterile saline for five days per week for two consecutive weeks under gaseous anesthesia (14, 26). Wild-type and *CD11c^PLXND1^ ^KO^* control mice were challenged with 35μl of sterile saline. The mice were sacrificed 48 hours after the final challenge to investigate the outcomes.

### Methacholine challenge test

Airway hyperresponsiveness (AHR) parameters, including airway resistance (Rn), tissue resistance (G), and tissue elastance (H), were evaluated using the FlexiVent animal ventilator (Scireq, Montreal, QC, Canada). Mice that received HDM or saline underwent thoracotomy, followed by intratracheal administration of an increasing gradient of methacholine dose (Saline,3, 6, 12, 25, and 50 mg/ml) at 5-minute intervals. Lung functions were investigated as previously described (14).

### Bronchoalveolar lavage fluid collection and differential cell count

Bronchoalveolar lavage fluid (BALF) was collected from the airways by two instillations of 1 ml sterile PBS containing 0.05 mM EDTA. Following centrifugation, the supernatant was stored at −80^◦C^ for future analysis.

Total BALF cells were counted using trypan blue and a hemocytometer. Cells were then prepared by cytospins, fixed, and stained with H&E. The differential cell count was blindly performed by two independent viewers, counting 200 cells, as previously reported (14).

### Immunophenotyping of BALF, spleen, blood, lymph node, and lung immune cells

Lungs, lymph nodes, blood, and spleen cells were utilized for immunophenotyping using FACS under the steady state condition. Tissues from *PLXND1^fl/fl^* (wild-type) and *CD11c^PLXND1^ ^KO^* mice were collected, and single-cell suspensions were prepared. Following washing and blocking with Fc-blocker, cells were stained with a mixture (0.5μl of antibodies/20μl of flow buffer per tube) containing the following anti-mouse antibodies using two antibody panels. The first panel consisted of fixable viability dye eFluor 780 (eBioscience), Siglec F-PE (clone E50-2440; BD Biosciences), CD11b-PE/Cy7 (clone M1/70), CD11c-PerCP/Cy5.5 (clone N418), Ly6G-allophycocyanin (clone 1A8), F4/80-FITC (clone BM8; all four from BioLegend). The second panel included fixable viability dye eFluor 780 (eBioscience), NK1.1-PE/Cy7 (clone PK136; eBioscience), CD3-PE (clone 145-2C11; eBioscience), CD4-allophycocyanin (clone GK1.5; Biolegend), and B220-FITC (clone RA3-6B2; BD Biosciences).

Moreover, inflammatory cells in the BALF were characterized using anti-mouse antibodies, including Siglec-F PE (clone E50-2440; BD Biosciences), CD11c Percp/Cy5.5 (clone N418), Ly6G-allophycocyanin (clone 1A8), CD11b PE/Cy7 (clone M1/70), F4/80 FITC (clone BM8; all four from BioLegend), and fixable viability dye APC-Cy7 (eBioscience). Subsequently, the samples were acquired using a BD FACSCanto II flow cytometer and analyzed using FlowJo software.

### Analyzing lung DC subsets and the expression of costimulatory molecules

Lungs were collected from *CD11c^PLXND1^ ^KO^* and *PLXND1^fl/fl^* mice challenged with either saline or HDM. The whole lung was minced and enzymatically digested in RPMI 1640 medium containing 1 mg/ml collagenase IV (Worthington Biochemical, Lakewood, NJ) at 37^◦C^ for 30 min. After red blood cells lysis with ACK (ammonium-chloride-potassium) buffer, the cells were counted and stained with anti-mouse antibodies (0.5μl of antibodies/20μl of flow buffer per tube) after Fc blocking. The antibody mixture included fixable viability dye eFluor 780 (eBioscience), CD45-eFluor 450 (clone 30F11), F4/80-FITC (clone BM8; eBioscience), anti-mouse CD11c-allophycocyanin (clone 418; eBioscience), MHC class II (I-A/I-E) eFluor 450 (clone M5/114.15.2; eBioscience), CD11b PE-Cy7 (clone M1/70; BioLegend), CD103 PerCP-Cy5.5 (clone 2E7; BioLegend), CD40-BV605 (clone 5C3; BioLegend), CD80-APC (clone 16-10A1; BioLegend), and CD86-APC-Cy7 (clone GL-1; BioLegend). Subsequently, the samples were acquired as described above.

### DC differentiation and isolation from bone marrow

Bone marrows were collected from naive *CD11c^PLXND1^ ^KO^* and *PLXND1^fl/fl^* mice. Red blood cells were depleted using ammonium chloride solution (ACK lysis buffer solution). Cells were cultured in RPMI 1640 (Sigma, St Louis, MO, USA) culture medium containing 10% FCS, 1% MEM non-essential amino acids solution (GIBCO, Berlin, Germany), 1% penicillin–streptomycin (GIBCO), 1% HEPES buffer solution (GIBCO), 1% sodium pyruvate (GIBCO), 50 lM 2-mercaptoethanol (GIBCO) and 20 ng/ml recombinant mouse granulocyte-macrophage colony-stimulating factor (GM-CSF) (PEPRO TECH EC LTD, London, UK) for 6 days at 37^◦C^ with 5% CO2 with medium changed every 2 days. Mature bone marrow-derived DC (mBMDCs) were obtained by overnight stimulation with 1mg/ml LPS (Sigma). Lastly, conventional DC were sorted using fixable viability dye APC-Cy7 (eBioscience), anti-mouse CD11c-allophycocyanin (clone 418; eBioscience), MHC class II (I-A/I-E) eFluor 450 (clone M5/114.15.2; eBioscience), using FACS flow cytometer. Purity of the cells was >92%, as checked by flow cytometry using a FACS Calibur (BD Biosciences, San Jose, CA) (27).

### B cell isolation and co-culture with BMDC

Spleen was harvested from a wild-type mouse, and single-cell suspention was prepared by depleting red blood cells using ammonium chloride solution (ACK lysis buffer solution). B cells were isolated using the B cell negative selection kit (EasySep™ Mouse B Cell Isolation Kit, Stem cell). The purity of the isolated B cells, as determined by flow cytometry using a FACS Calibur instrument, was consistently above 95%.

For the co-culture experiment, 50 μl of B cells at a concentration of 2 x 10^6^ cells/ml, and 50 μl of BMDC at 2 x 10^6^ cells/ml (1:1 ratio) were co-cultured for 4 days. Cells were stimulated with 0.5ug/ml purified NA/LE hamster anti-mouse-CD40 (clone HM40-3 BD Pharmingen), 10ug/ml affinity pure F(ab)2 fragment goat anti-mouse-IgM, (clone 115-006-020; Jackson ImmunoResearch), and 25ng/ml recombinant mouse IL-4 (clone 404-ML-025/CF; R&D System) in final volume of 200 μl of RPMI 1640 medium supplemented with 10% FBS, 2 mM L-glutamine, 100 U/ml penicillin,100 U/ml streptomycin, and 50 mM 2-ME, in 96-well plates, and incubated at 37^◦C^ with 5% CO2. After four days, the supernatant was collected, and IgE production was measured using ELISA (27).

### Cytokine measurement

Mesoscale ELISA was used to assess IL-4, IL-5, IL-13, IL-17A, IFN-γ, and MCP-1 levels in BALF supernatants according to the manufacturer’s instructions. ELISA data were analyzed using SoftMax Pro software (Molecular Devices). All cytokine ELISA kits were purchased from BioLegend (San Diego, CA), except for IL-13 (eBioscience).

### Intracellular cytokine detection

The intracellular staining of cytokines was conducted as previously described (14). Briefly, mediastinal lymph node cells were cultured for 4 hours at 37°C with 5% CO2 and stimulated with PMA, ionomycin, and the protein transport inhibitor brefeldin A (Invitrogen).

Cells were then collected, and extracellular staining was performed using anti-mouse CD3 PE/Cy7 (clone 145-2C11; eBioscience) and CD4-allophycocyanin (clone G1.5; eBioscience). Subsequently, intracellular staining was performed using specific anti-mouse antibodies, including IFN-γ PerCP-Cy5.5 (clone XMG1.2; eBioscience), IL-4 PE (clone 11B11; eBioscience), and IL-17A (clone TC11-18H10.1; BioLegend). The samples were acquired using the FACSCanto II flow cytometer and analyzed with FlowJo software.

### Measurement of Immunoglobulins in serum

Serum samples were obtained from *CD11c^PLXND1^ ^KO^*and *PLXND1^fl/fl^* mice that received either saline or HDM. The total and HDM-specific IgE and IgG1 were measured using ELISA, according to the manufacturer’s instructions (14, 28). ELISA antibodies for measuring total and HDM-specific Igs in serum were purchased from Southern Biotech (Birmingham, AL). ELISA data were analyzed using SoftMax Pro software (Molecular Devices).

### Lung histology

The left lobe of the lung was dissected and fixed in formalin overnight, followed by embedding in paraffin. The severity of airway inflammation, mucus production, and collagen deposition in lung tissue sections were investigated by performing H&E, periodic acid Schiff (PAS), and Sirius red staining, respectively in *CD11c^PLXND1^ ^KO^*and wild-type mice following saline or HDM administration (14, 29). The staining results were evaluated by a pathologist, and reported as a pathological score.

### Real-time PCR

Total RNA was isolated from the middle lobe of the lung using TRIzol (Ambion). MultiScribe reverse transcriptase was performed for 1mg of RNA to synthesize cDNA according to the manufacturer’s instructions (Applied Biosystems, Foster City, CA). The expression of the collagen (*COL3*), and mucin (*MUC5AC*) genes was analyzed by quantitative real-time PCR (qRT-PCR) (Table1). Eukaryotic elongation factor 2 (EEF2) was used as a housekeeping gene. qRT-PCR was done in a 96-well optical plate with an initial one-cycle denaturation step for 10 min at 95◦C, 40 cycles of PCR (95^◦C^ for 15s, 60^◦C^ for 30s, and 72^◦C^ for 30s), one cycle of melting, and one cooling cycle (Bio-Rad CFX96 real-time PCR system). Product specificity was assessed by performing a melting curve analysis and examining the quality of amplification curves. The amplification of target genes was calculated by normalizing by the amplification of EEF2 (Δ^Ct^) and then normalizing by control groups (ΔΔ^Ct^) (30).

**Table1.**
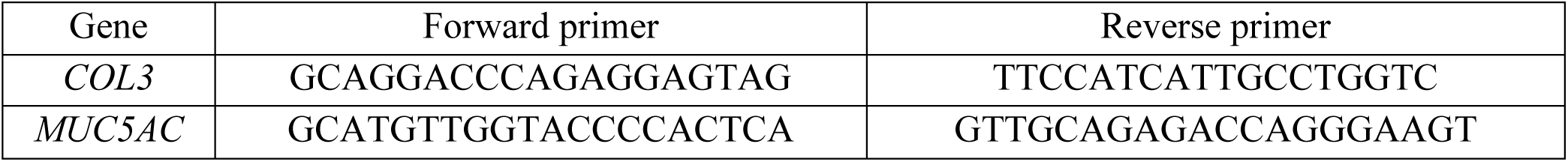
The forward and reverse primers are used to investigate gene expression.

### Statistical analyses

GraphPad Prism 9.0 software was used for statistical analysis. Depending on the number of groups and treatments, data were analyzed by an unpaired t-test, one-way ANOVA, or two-way ANOVA, followed by a Tukey test. Differences were statistically significant at *p < 0.05, **p < 0.01, and ***p < 0.001.

## Results

### PLXND1 ablation in the CD11c+ DC did not affect immune cells composition at the steady state

To investigate whether the absence of *PLXND1* in CD11c+ DC alters immune cell composition, we conducted FACS-based immunophenotyping of various immune cell populations, including neutrophils, eosinophils, alveolar macrophages (AM), interstitial macrophages (IM), NK cells, B cells, and T cells, in the lungs, spleen, mediastinal lymph nodes, and blood of *CD11c^PLXND1^ ^KO^*and *PLXND1^fl/fl^* mice (wild type).

Our finding revealed that at baseline, there were no significant differences in the number of these immune cell populations between *CD11c^PLXND1^ ^KO^* and *PLXND1^fl/fl^* mice in the lungs, spleen, lymph nodes, and blood (Figure 1D, 1E, 1F, and 1G). These results indicate that the absence of *PLXND1,* specifically in CD11c+ DC, does not impact immune cell composition in different tissues under steady-state conditions.

**Figure 1.**
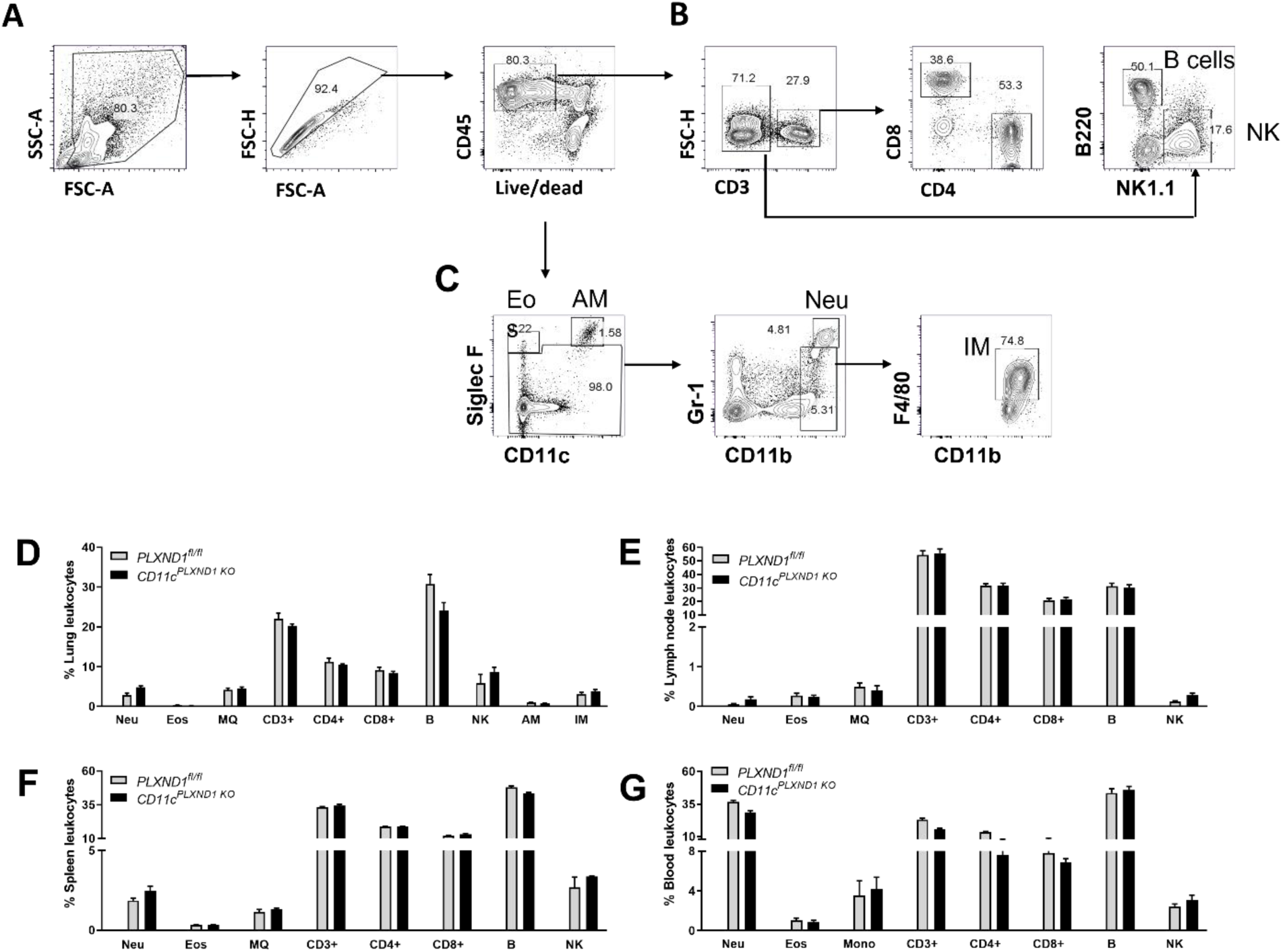
Immunophenotyping of inflammatory cells in *CD11c^PLXND1 KO^* and *PLXND1^fl/fl^* mice. Lungs, spleen, blood, and lymph nodes were harvested from CD11cPLXND1 KO and wild-type mice. Following enzymatic digestion, single-cell suspensions were analyzed using FACS with specific antibodies to characterize various inflammatory cell populations at the steady state. **(A)** The general gating strategy involved excluding debris and doublet cells, followed by selecting viable leucocytes from the total cell population. **(B)** T and B cells were characterized based on the surface expression of CD3 and B220, respectively, followed by further identification of CD4+ and CD8+ cells within the CD3-expressing cell population. Pulmonary NK cells were identified as NK1.1+ cells. **(C)** Eosinophils were characterized by expressing Siglec-F+/CD11c-, and alveolar macrophages by expressing Siglec-F+/CD11c+. Neutrophils were identified by the expression of Ly6G+ (1A8) and CD11b+. Interstitial macrophages were identified as F4/80+/CD11b+ cells. The numbers of each cell type in **(D)** lung, **(E)** mediastinal lymph node, **(F)** spleen, and **(G)** blood were compared between CD11cPLXND1 KO and WT mice under the steady-state condition. Data are presented as the mean (pre-gated on CD45+) with standard error of the mean (SEM). All data shown are representative of four to five mice per group. Eos: Eosinophil; Neu: Neutrophil; AM: Alveolar macrophage; IM: Interstitial macrophage; MONO: Monocyte; NK: Natural killer cell.

### The PLXND1 deficiency in CD11c+ DC exaggerated airway hyperresponsiveness upon HDM exposure

To assess the impact of *PLXND1* ablation in CD11c+ cells on airway hyperresponsiveness (AHR), both *CD11c^PLXND1^ ^KO^*and *PLXND1^fl/fl^* (wild type) mice were subjected to HDM allergen for five consecutive days over two weeks (Figure 2A) (31, 32). A significant increase was observed in airway resistance (Rn) and tissue elastance (H) in *CD11c^PLXND1^ ^KO^* mice compared to *PLXND1^fl/fl^*mice (Figure 2B and 2D). However, no difference was detected in tissue resistance (G) (Figure 2C). These findings indicate that the absence of *PLXND1* in CD11c+ DCs exacerbates HDM-induced airway hyperresponsiveness in the acute HDM model of allergic asthma.

**Figure 2.**
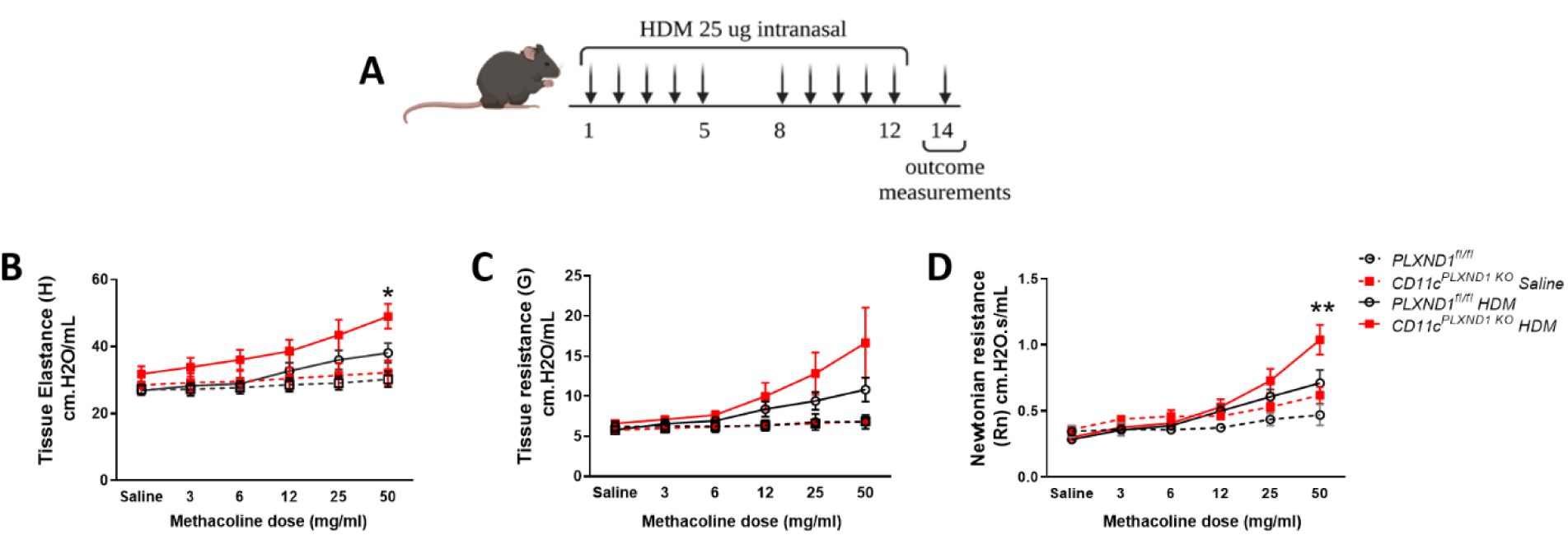
Mice with *PLXND1* deletion in CD11c+ DC exhibited enhanced airway hyperresponsiveness. **(A)** The allergic airway disease model was established by intranasal exposure to HDM for two weeks, while control mice received saline. *CD11c^PLXND1^ ^KO^* and WT mice underwent a tracheotomy and were subsequently challenged with methacholine to measure **(B)** airway resistance, **(C)** tissue resistance, and **(D)** tissue elastance. All data presented are representative of three to five mice per group; 2-Way ANOVA, *p<0.05 **p<0.01.

### The PLXND1 deficiency in CD11c+ DC enhanced airway inflammation upon HDM exposure

To assess the impact of *PLXND1* ablation in CD11c+ cells on airway inflammation and cell recruitment into the lungs, we investigated the total number of cells in bronchoalveolar lavage fluid (BALF), and the presence of neutrophils, eosinophils, and macrophages, using flow cytometry (14). We found that the numbers of BALF total cells, eosinophils, neutrophils, and alveolar macrophages were not significantly altered in the airways of HDM-treated *CD11c^PLXND1^ ^KO^* mice compared to *PLXND1^fl/fl^* mice (Figure 3A, 3B, 3C, 3D). However, the number of interstitial macrophages significantly increased in *CD11c^PLXND1^ ^KO^* compared with *PLXND1^fl/fl^* mice (Figure 3E).

**Figure 3.**
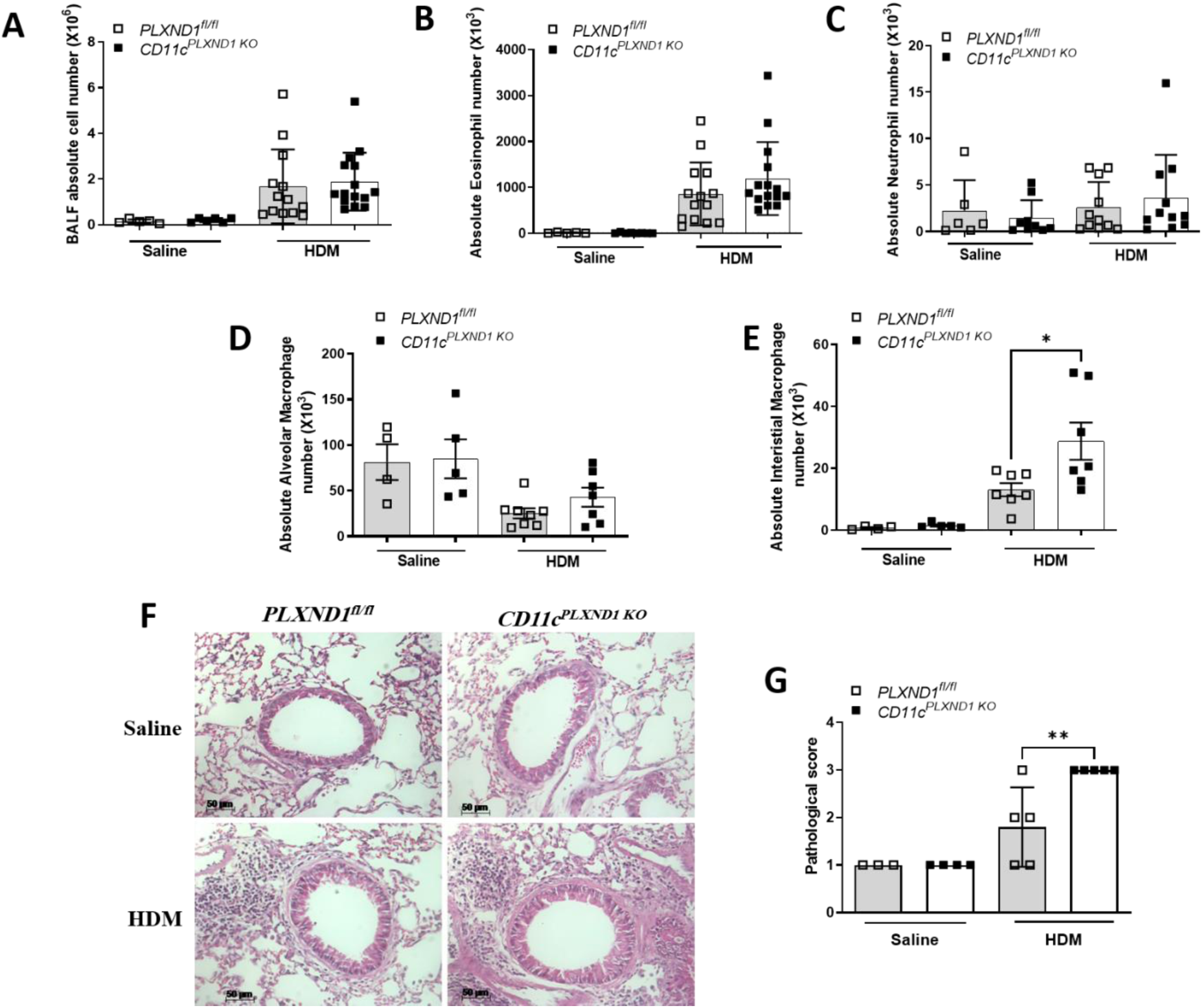
Mice with *PLXND1* ablation in CD11c+ DC showed increased lung inflammation. **(A)** The total number of BALF cells, **(B)** eosinophils, **(C)** neutrophils, **(D)** alveolar macrophages, and **(E)** interstitial macrophages in the BALF were investigated using flow cytometry. **(F)** Lung inflammation was evaluated by performing H&E staining, and **(G)** the results were reported as a pathological score. Scale bars, 50um; All data presented are representative of three to five mice per group; 2-Way ANOVA, *p<0.05 **p<0.01.

In contrast to the BALF, we demonstrated the increased recruitment of inflammatory cells in the lungs of *CD11c^PLXND1^ ^KO^* mice compared to *PLXND1^fl/fl^* mice using H&E staining of lung tissue sections (Figure 3F & G). These findings collectively indicate that, following selective deletion of *PLXND1* in CD11c+ DCs, inflammation is significantly enhanced during asthma, as indicated by the recruitment of inflammatory cells and overall histopathological changes in lung tissue.

### Lack of PLXND1 in CD11c+ DC exaggerated goblet cell hyperplasia and collagen3 gene expression upon HDM exposure

Given that goblet cell proliferation and sub-epithelial fibrosis contribute to airway remodeling in allergic asthma (33), we investigated whether the absence of *PLXIND1* in CD11c+ DC affects mucin production and collagen deposition. To visualize mucus production and collagen deposition, we performed Sirius red and periodic acid-Schiff staining on lung tissue sections, respectively. Our results revealed significantly higher mucus but not collagen deposition in the *CD11c^PLXND1^ ^KO^* mice compared with *PLXND1^fl/fl^* mice (Figure 4A and 4C).

**Figure 4.**
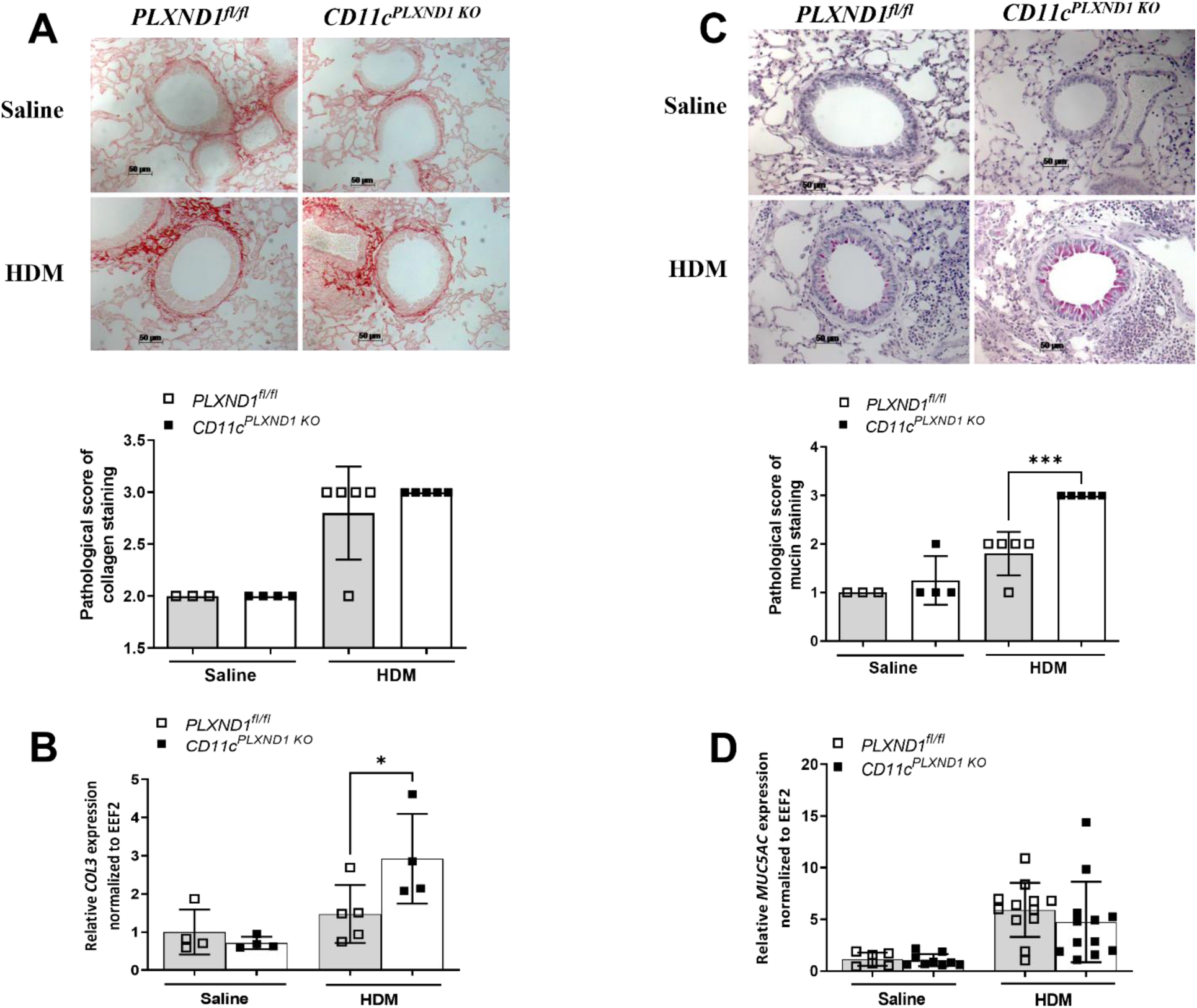
*PLXND1* deletion in CD11c+ DC enhanced changes in structural cells and airway remodeling. **(A)** Collagen deposition and **(C)** Mucus production were visualized using Sirius red and periodic acid-Schiff (PAS) staining respectively. **(B)** Collagen 3 (*COL3*) and **(D)** *MUC5AC* expression were assessed by real-time PCR in *CD11c^PLXND1^ ^KO^* and *PLXND1^fl/fl^* mice. . Scale bars, 50um; All data presented represent three to five mice per group; 2-Way ANOVA, *p<0.05 ***p<0.001.

To validate these findings, we investigated the expression of the *COL3* and *Muc5AC* genes in the lung tissue. HDM-subjected *CD11c^PLXND1^ ^KO^* mice exhibited significantly higher *COL3* gene expression compared to *PLXND1^fl/fl^* mice (Figure 4B). However, no significant difference was found in the *Muc5AC* gene expression between the two groups (Figure 4D).

These findings indicate that the ablation of *PLXND1* in CD11c+ DCs leads to increased airway remodeling events during allergic asthma, as evidenced by enhanced mucus production, higher *COL3* gene expression, and subsequent deposition.

### Ablation of PLXND1 in CD11c+ DC affects the production of CCL2/MCP1

Various cytokines, particularly Th2/Th17 cytokines, play critical roles in the induction and exacerbation of inflammation and remodeling during allergic asthma (34). Accordingly, we investigated the levels of inflammatory cytokines in the BALF of *CD11c^PLXND1^ ^KO^* and *PLXND1^fl/fl^* mice. Although there is a slight increase in the levels of cytokines in the BALF of *CD11c^PLXND1^ ^KO^*, the deletion of *PLXND1* in CD11c+ DC did not statistically affect the production of IL-4, IL-5, IL-13, IL-17A, and IFN-γ in the BALF of *CD11c^PLXND1^ ^KO^* mice compared to *PLXND1^fl/fl^* mice upon HDM challenge (Figure 5A, 5B, 5C, 5D, 5E). However, the levels of CCL2/MCP-1 (Monocyte chemoattractant protein-1) were enhanced in *CD11c^PLXND1^ ^KO^* mice compared with *PLXND1^fl/fl^* mice (Figure 5F).

**Figure 5.**
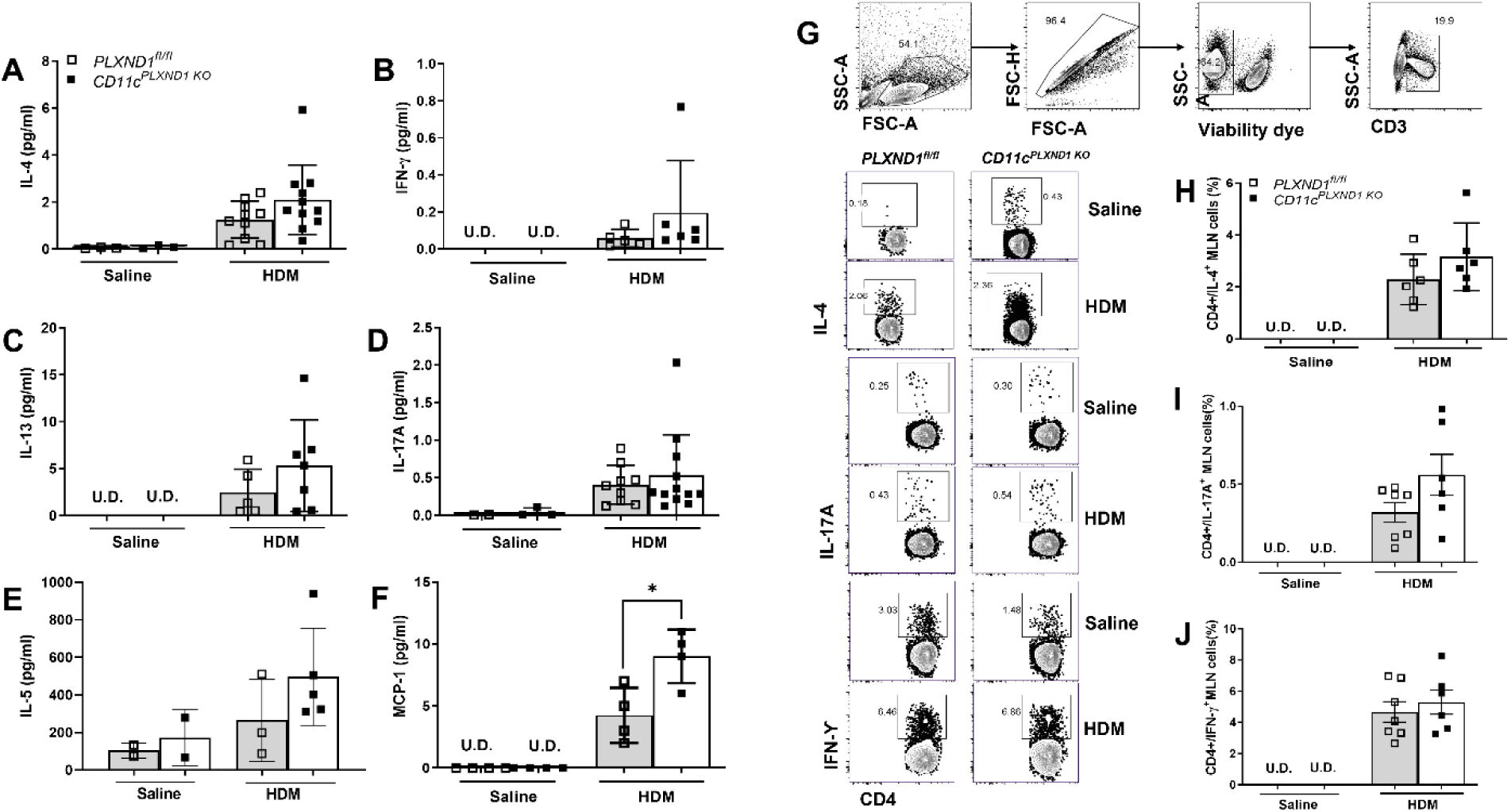
Lack of *PLXND1* in CD11c+ DC affects the production of CCL2/MCP1 in allergic asthma. The levels of **(A)** IL-4, **(B)** IFN-γ, **(C)** IL-13, **(D)** IL-17A, **(E)** IL-5, and **(F)** MCP-1 were measured by mesoscale ELISA in BALF supernatants obtained from *CD11c^PLXND1^ ^KO^* and WT mice after intranasal exposure to saline or HDM. FACS analysis using specific antibodies was performed on lymph node cells of *CD11c^PLXND1^ ^KO^* and WT mice to characterize T cell-derived cytokines. **(G)** The general gating strategy selected lymphocytes and single cells, followed by including viable and CD3+ cells as T lymphocytes. T cell-derived cytokine production was assessed by gating on CD4+ cells and the levels of **(H)** IL-4, **(I)** IL-17A, and **(J)** IFN-γ were determined. Data are presented as mean with SEM. All data are representative of three to five mice per group. 2-Way-ANOVA, *p<0.05

Moreover, to further assess the T cell subsets and their production of specific cytokines, mediastinal lymph node (MLN) were harvested from *CD11c^PLXND1^ ^KO^*and *PLXND1^fl/fl^* mice, and MLN cells were stimulated with stimulation cocktail *ex vivo* and analyzed using surface and intracellular markers by flow cytometry, as previously described (35). IL-4, IL-17A, and IFN-γ levels did not change after stimulation of the MLN cells from *CD11c^PLXND1^ ^KO^* compared to WT mice (Figure 5H, 5I, and 5J).

These results suggest that except for CCL2/MCP1, the lack of *PLXND1* in CD11c+ DC does not impact the biosynthesis and release of IL-4, IL-5, IL-13, IL-17A, and IFN-γ in this HDM model of allergic asthma.

### Total and HDM-specific serum IgE levels were enhanced in response to the ablation of PLXND1 in CD11c+ DC

We measured the production of total and HDM-specific Igs in serum isolated from *CD11c^PLXND1^ ^KO^* and *PLXND1^fl/fl^*mice using ELISA. The total and HDM-specific IgE levels significantly increased in *CD11c^PLXND1^ ^KO^* mice compared to *PLXND1^fl/fl^* mice upon HDM challenge (Figure 6A and 6B). However, no difference was detected in total and HDM-specific IgG1 levels following HDM exposure (Figure 6C and 6D). These results suggest that the lack of *PLXND1* on CD11c+ DC enhances IgE levels, which may contribute to exacerbated allergic reaction in asthma.

**Figure 6.**
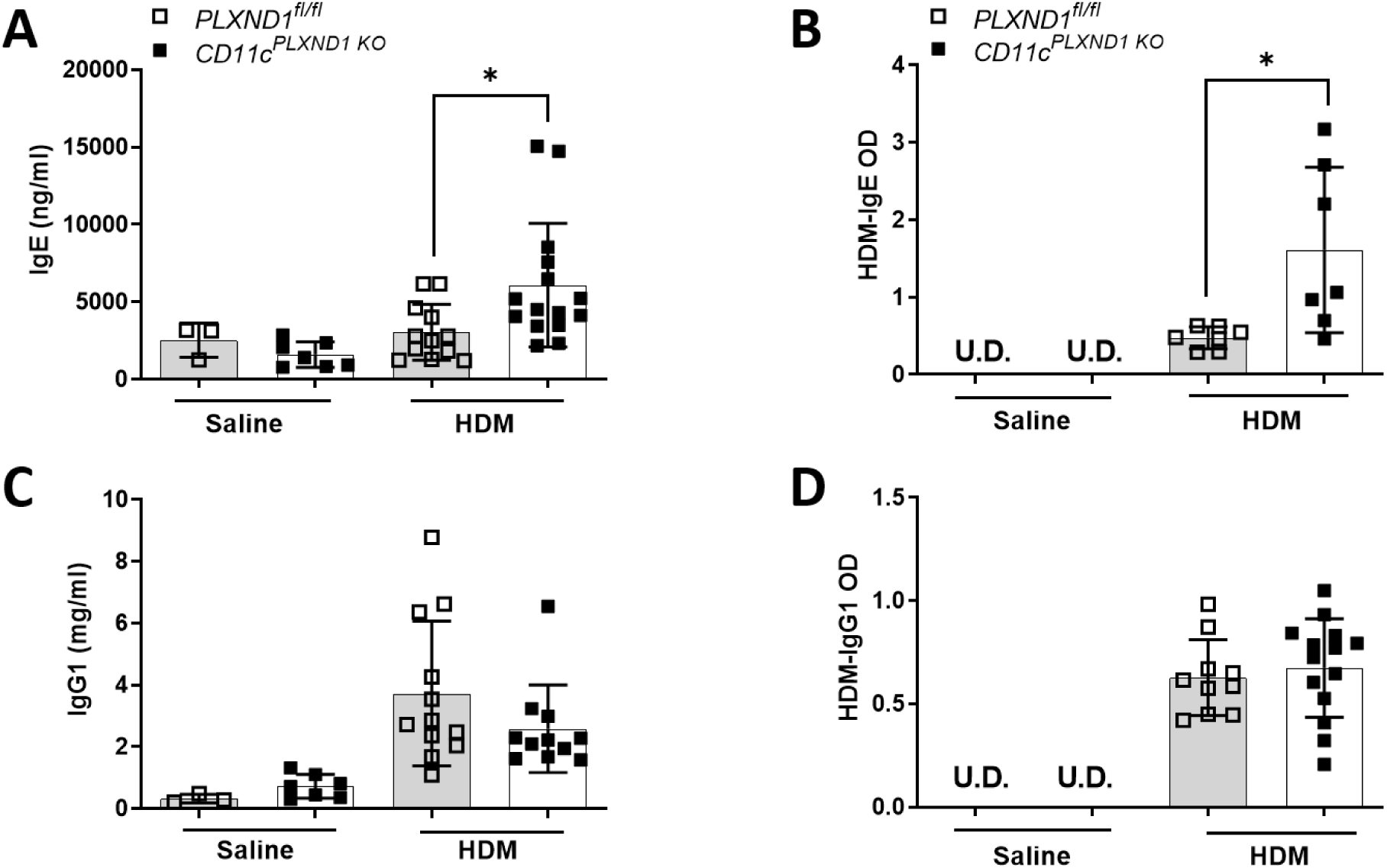
Lack of *PLXND1* in CD11c+ DC increased IgE levels in allergic asthma. After intranasal sensitization and challenge with either saline or HDM, serum samples were collected from *CD11c^PLXND1^ ^KO^* and WT mice. Then the levels of **(A)** total and **(B)** HDM-specific IgE, as well as **(C)** total and **(D)** HDM-specific IgG1, were measured using ELISA. Data are presented as mean with SEM. All data are representative of three to five mice per group. 2-Way ANOVA *p<0.05.

### Deletion of PLXND1 in CD11c+ DC leads to an increase in the recruitment of cDC2s into the lungs

Lung DCs comprise heterogeneous populations, such as conventional DCs categorized into CD11b+ (cDC2) and CD103+ (cDC1) subtypes. CD11b+ cDC2 cells are involved in Th2/Th17 priming and exacerbate atopic responses in the airways, while CD103+ cDC1 cells act as tolerogenic cells and ameliorate inflammation following exposure to HDM (36).

We observed a significant increase in the number of cDCs (CD11c+/MHCII+) in the airways of *CD11c^PLXND1^ ^KO^* mice compared with *PLXND1^fl/fl^*mice (Figure 7B). Additionally, the number of CD11b+ cDC2 was significantly higher compared to CD103+ cDC1 upon HDM challenge in *CD11c^PLXND1^ ^KO^* mice (Figure 7C, 7D).

**Figure 7.**
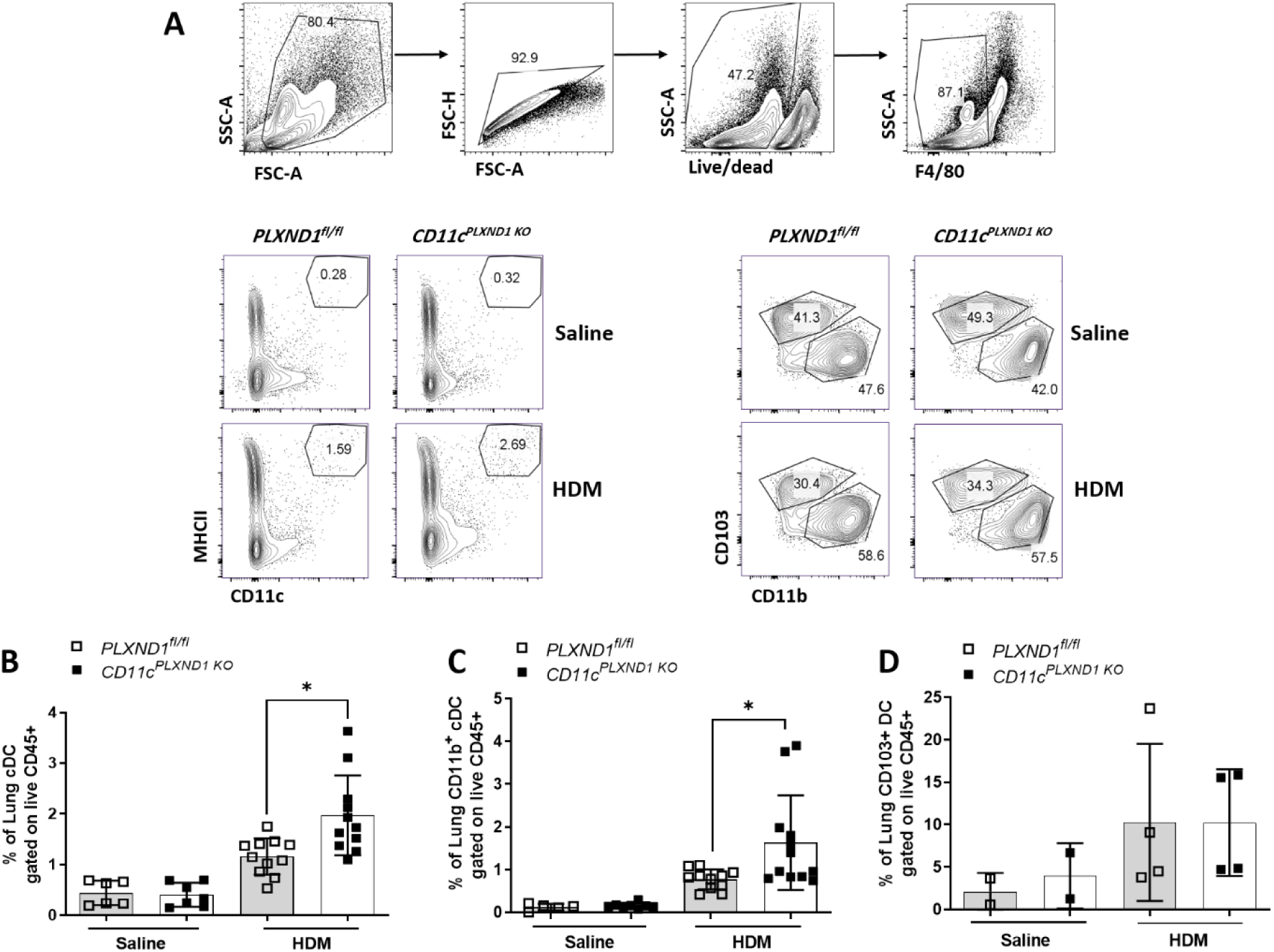
*PLXND1* deletion in CD11c+ DC increased the cDCs recruitment into the lungs. Lungs were harvested from *CD11c^PLXND1^* ^KO^ and WT mice after intranasal exposure to saline or HDM. **(A)** Pulmonary DC subsets were characterized using specific antibodies for flow cytometry. After excluding doublets and dead cells, macrophages were excluded by gating on F4/80 negative cells. **(B)** The number of pulmonary DC (MHCII^hi^/CD11c+) and the subsets of **(C)** CD11b+ and **(D)** CD103+ DC were compared between *CD11c^PLXND1^ ^KO^* and *PLXND1^fl/fl^* mice. Data are presented as the mean (pre-gated on CD45+) with standard error of the mean (SEM). Pulmonary DC data represent four independent experiments generated from four pooled whole-lung samples in each group per experiment. All data are representative of three to five mice per group. 2-Way ANOVA, *p<0.05.

Collectively, these findings demonstrated that lack of *PLXND1* in DC increased the recruitment of pulmonary cDCs mostly CD11b+ DC subtype, which was further associated with enhanced CCL-2/MCP-1 levels and worsened allergic asthma features.

### Deletion of PLXND1 in CD11c+ DC enhanced the IgE levels ex vivo

To further assess the impact of *PLXND1* deficiency in CD11c+ DC on B cell function and IgE production, conventional DCs were differentiated from bone marrow of *CD11c^PLXND1^ ^KO^* and *PLXND1^fl/fl^* mice, and co-cultured with isolated splenic B cells from WT mice *ex vivo* (Figure 8A). Notably, the levels of IgE were significantly higher in *CD11c^PLXND1^ ^KO^*-isolated DC-B cell co-culture compared to DC isolated from *PLXND1^fl/fl^* mice (Figure 8B).

**Figure 8.**
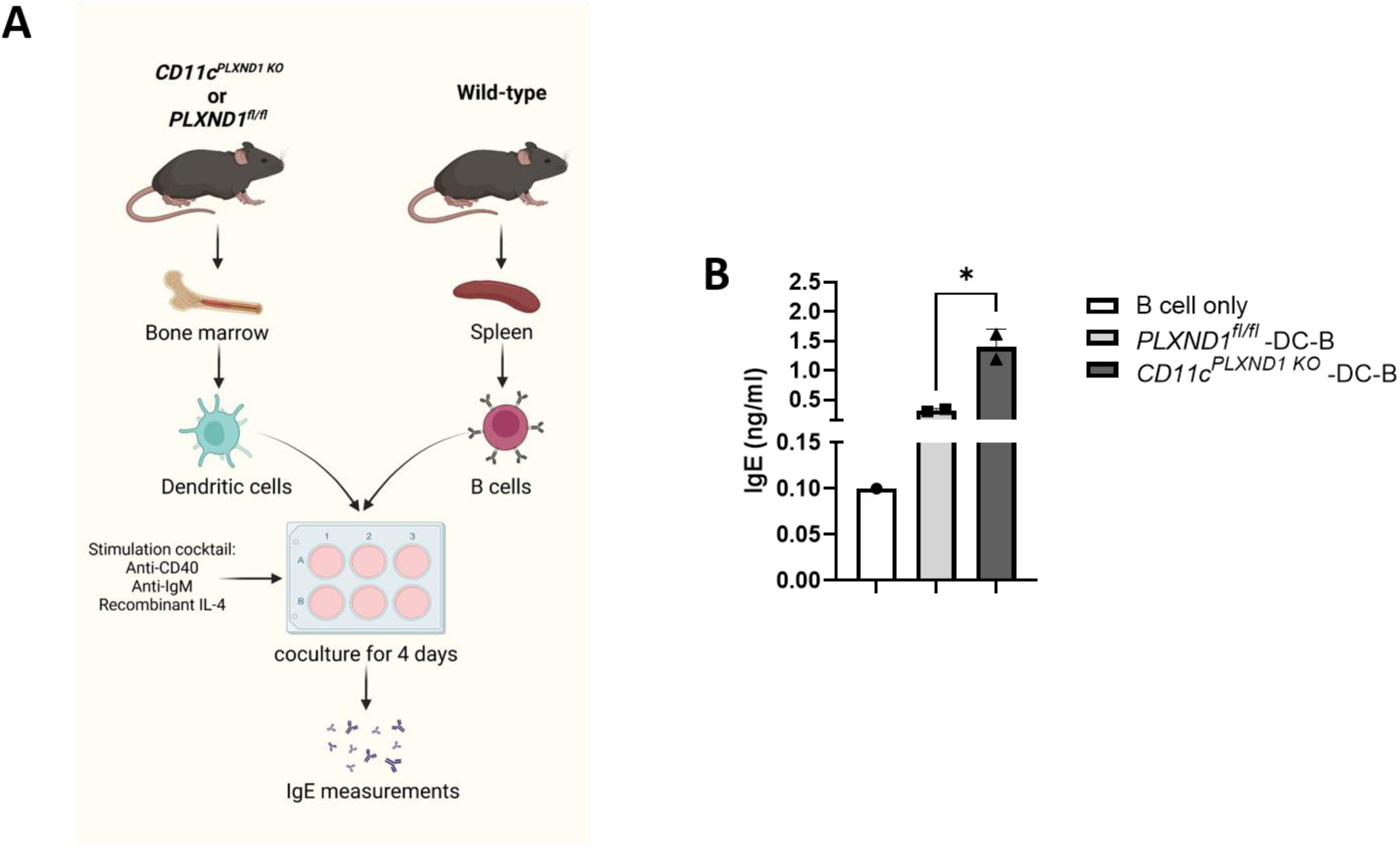
Lack of *PLXND1* in CD11c+ DC induced IgE production *ex vivo*. **(A & B)** To further investigate the impact of plexinD1 deficiency in CD11c+ DC, bone marrow-derived DCs from *CD11c^PLXND1^ ^KO^* and *PLXND1^fl/fl^* mice co-cultured with splenic B cells isolated from WT mice to assess the IgE levels *ex vivo*. The supernatant of DC-B cell co-culture were collected and IgE levels were measured using ELISA. All data are representative of two to three mice per group. Data are presented as mean with SEM. 1-Way ANOVA, *p<0.05.

Moreover, considering the crucial role of costimulatory molecules in the initiation of immune responses and Ig class switching by DCs (37, 38), we further assessed the level of costimulatory molecules in splenic CD103+ cDC1 and CD11b+ cDC2. However, no difference in the expression of costimulatory molecules, including CD40, CD80, CD86, BAFF, APRIL, and PDL-1, was observed upon HDM challenge between *CD11c^PLXND1^ ^KO^* and *PLXND1^fl/fl^*mice (data not shown). As *Eda K. Holl et al.* showed absence of *PLXND1* in DCs did not affect the ability of these cells to upregulate costimulatory molecules (39). These findings suggested that the lack of *PLXND1* in DC enhanced IgE levels by B cells, which further explains the exacerbated allergic asthma reactions observed in our model.

## Discussion

In this study, we addressed the impact of *PLXND1* deficiency in CD11c+ DC in an HDM allergic model of asthma (40). We demonstrated that the lack of *PLXND1* in CD11c+ DC exacerbates airway hyperresponsiveness (AHR) parameters, such as airway resistance and tissue elastance. We also observed higher mucus production and an increase in the collagen gene expression in the lungs of *CD11c^PLXND1^ ^KO^* DC compared to WT counterparts suggesting the role of CD11c+ DC *PLXND1* deficiency in airway remodelling. Moreover, we found higher airway inflammation, particularly an enhanced number of interstitial macrophages and elevated CCL-2/MCP-1 levels in the BALF of *CD11c^PLXND1^ ^KO^* DC mice. The absence of *PLXND1* in CD11c+ DC resulted in higher total and HDM-specific serum IgE levels and enhanced recruitment of CD11b+ cDC2 to the lungs. Mechanistically, co-culture of B cells with DC from *CD11c^PLXND1^ ^KO^* DC mice led to significantly higher IgE levels *ex vivo* compared to the DC isolated from WT mice. Our data highlighted that the Sema3E-plexinD1 axis in CD11c+ DC is critical in modulating allergic asthma features, including AHR, remodelling and inflammation.

DCs play a central role in recognizing allergens in the airways and producing IgE antibodies, thereby triggering subsequent allergic responses in individuals with IgE-exacerbated asthma (41). CD11b+ DC, cDC2, are essential in initiating and shaping the immune response, including the production of IgE antibodies from activated B cells (41). DCs by capturing and processing allergens, prime the immune system to recognize and respond to potential stimuli, like HDM (42). Once encountered, DCs migrate to mediastinal lymph nodes, where they present allergens to naive T cells (42). Crucially, DCs promote the differentiation of naive T cells into Th2 cells, a subset specialized in orchestrating allergic reactions (42). Through the release of specific cytokines, like IL-4 and IL-13, DCs drive the activation and proliferation of Th2 cells, which in turn stimulate B cells to differentiate into IgE-producing plasma cells (42). The resulting IgE antibodies circulate in the bloodstream and tissues, awaiting re-exposure to the same allergen (43). Upon subsequent encounters, IgE binds to mast cells and basophils, releasing inflammatory mediators like histamine and leukotrienes (43). This cascade of events leads to airway inflammation, smooth muscle contraction, mucus production, and ultimately bronchoconstriction, precipitating asthma symptoms (43). PlexinD1 is highly expressed in lung and bone marrow-derived DCs and mature and immature DCs (17, 39, 44). Notably, plexinD1 is involved in B cells homing into germinal centers, induction of humoral responses, regulation of long-lived bone marrow plasmacytes, and recall humoral memory responses (45).

In this study, we observed that ablation of *PLXND1* in CD11c+ DC increased the total number of cDCs in the lungs and further elevated the number of CD11b+ cDC2 compared to CD103+ cDC1. These data are consistent with our prior studies, which showed a higher number of CD11b+ DCs than CD103+ DCs in the airways of *Sema3E knockout (KO)* mice at the baseline and upon HDM sensitization (17, 18). Moreover, these CD11b+ DC demonstrated more migration and HDM uptake in our *Sema3E KO* allergic asthma model (17). CD11b+ DCs are associated with Th2/Th17 immunity, driving IgE production, while CD103+ DCs promote Th1 responses and induce tolerance, inducing IgA production in response to inhaled allergens (46–48) In line with this, we observed elevated total and HDM-specific IgE levels in the serum of *CD11c+ PLXND1 KO* mice compared to wild-type mice. Co-culturing BMDCs from *CD11c+ PLXND1 KO* mice with B cells from wild-type mice resulted in higher levels of IgE *ex vivo* compared to DCs isolated from wild-type mice. These findings agree with the effect of Sema3E deficiency and *PLXND1* deficiency in Cx3cr1 interstitial macrophages on IgE production (14, 35). However, the exact mechanism by which plexinD1 in DCs can affect IgE production needs further investigation. Collectively, our data suggest that the absence of the plexinD1-Sema3E axis in CD11c+ DC can regulate IgE production by the B cells and exacerbate allergic asthma via an IgE-mediated mechanism.

Airway epithelial cells (AECs) have the potential to regulate DCs and macrophages in the lungs by expressing a variety of molecules that can modulate their behavior either positively or negatively through direct and indirect mechanisms (49, 50). AECs are a significant source of MCP-1/CCL-2 within the lungs, as observed in human asthmatic patients subjected to allergen challenges (51). MCP-1 plays a pivotal role in the infiltration and migration of monocytes/macrophages to the lungs (51, 52). In this study, elevated levels of MCP-1 were observed in the BALF of *CD11c^PLXND1^ ^KO^* DC mice, accompanied by an increased number of interstitial macrophages and conventional type-2 DCs compared to their wild-type counterparts.

TNF can stimulate the production of MCP-1 by epithelial cells via the MAPK signalling pathway (53). TNF can be produced by DCs and macrophages during the initial stages of immune responses (54, 55). Furthermore, our prior study has shown that plexinD1 on macrophages modulates TNF production through the MAPK, STAT, and NF-κB signaling pathways (56). In the current study, it is plausible to hypothesize that plexinD1 deficient CD11c+ DCs induces the production of MCP-1 by airway epithelial cells, subsequently promoting the infiltration of interstitial macrophages into lung tissue in a TNF-dependent manner. However, additional investigations are necessary to comprehensively understand the mechanisms through which plexinD1 in CD11c+ DCs regulates the production of MCP-1 by airway epithelial cells. In summary, our study underscores the influence of the Sema3E-plexinD1 complex on CD11c+ DCs in shaping airway inflammation during allergic asthma.

Airway hyperresponsiveness (AHR) is the most characteristic clinical feature of asthma, primarily induced by airway inflammation (57); activated pulmonary DCs trigger the latter through the induction of a Th2/Th17 response (58). In this study, we demonstrated that the absence of *PLXND1* in CD11c+ DCs exacerbated bronchial hyperreactivity, including airway resistance (Rn) and tissue elastance (H). These findings align with our previous study, where *PLXND1* deletion in Cx3cr1 interstitial macrophages resulted in aggravated airway resistance (Rn) (35). Furthermore, we previously revealed worsened AHR parameters, such as airway resistance (Rn), tissue resistance (G), and tissue elastance (H), in allergic asthma with a global absence of Sema3E, the canonical ligand for plexinD1 (13–16, 18, 19).

Allergen encounter leads to significant recruitment of DCs into the airways (59). These DCs activate various subtypes of T cells, leading to enhanced recruitment of immune cells and production of proinflammatory cytokines, such as IL-4, IL-5, IL-9, IL-13, IL-17A, and IFN-γ (58). These cytokines can trigger smooth muscle cells and induce AHR (34, 58). Interestingly, specific elimination of conventional DCs can prevent allergic airway inflammation and AHR (60, 61). As observed in this study, the number of conventional DCs, particularly cDC2s, responsible for inducing type 2 inflammation, increased in the lungs in response to *PLXND1* deletion in CD11c+ DC. Although type 2 cytokine levels showed no difference, we observed a significantly higher level of MCP-1 in our CD11c+ DC *PLXND1* KO model, which led us to speculate that the ablation of *PLXND1* in DCs by increasing inflammation, induced AHR in our model. MCP-1 has been shown to induce AHR by directly activating and degranulating mast cells (62, 63). Notably, Anti-MCP-1 antibodies inhibited methacholine-induced AHR and reduced histamine release into the BALF of the cockroach-induced allergic model (62). Altogether, the regulated recruitment of DC subsets or modulation of their functions by Sema3E-plexinD1 may be linked to the increased AHR observed in our CD11c+ DC *PLXND1* KO model. Further studies are needed to understand how plexinD1 in the CD11c+ DC regulates airway resistance and tissue elastance.

Collagen deposition is considered an essential aspect of asthma (64), and along with the excessive mucin production by goblet cells, it reduces the radius of the airways, restraining airflow and resulting in airway resistance in asthma (65). We demonstrated that the ablation of *PLXND1* in CD11c+ DC enhanced collagen gene expression, and a significant increase in mucus production in the airways was observed. These findings align with our previous studies, where the deletion of *PLXND1* in Cx3cr1 interstitial macrophages (35) and the global absence of Sema3E resulted in increased collagen deposition in the lamina reticularis, the expression of *Muc5AC* and *Muc5b*, and hypersecretion of mucus in the airways (14, 15, 35).

The exact mechanism of how Sema3E-plexinD1 participates in fibrosis during asthma remains unclear. However, DCs, through interactions with other inflammatory cell types and their mediators, can contribute to airway remodeling (66). In this study, the number of interstitial macrophages and the levels of MCP-1 significantly increased in response to the deletion of *PLXND1* in CD11c+ DC. MCP-1 stimulates fibroblasts, the cells responsible for producing collagen and other extracellular matrix components, leading to increased production and deposition of collagen (67); thus contributing to the fibrotic remodelling of lung tissue (63). Furthermore, MCP-1 has been shown to induce the differentiation of blood-recruited fibroblasts into myofibroblasts (68), contributing to tissue scarring and fibrosis. Lastly, MCP-1 can interact with other pro-fibrotic factors, such as transforming growth factor-beta (TGF-β), to promote fibrosis synergistically (67). TGF-β is a potent inducer of fibrosis, and its effects can be potentiated by MCP-1 (67). MCP-1 has been found to interact with other proinflammatory cytokines, such as interleukin-13 (IL-13) (69, 70) and activates p44/42MAPK, a kinase that plays a crucial role in mucin regulation in the bronchial epithelium (71).

Overall, it is enticing to suggest that the impact of CD11c+ DC *PLXND1* KO on airway remodelling and fibrosis could be mediated by direct interaction of MCP-1 with fibro/myofibroblast or indirectly by increased recruitment of interstitial macrophages (i.e. M2 macrophages) into the lung and their associated-pulmonary fibrosis (72). Sema3E-plexinD1 signalling in CD11c+ DC is critical for remodelling events during asthma, as evidenced by higher mucus and collagen in the airways and increased stiffness or tissue elastance in our model.

In conclusion, our data show that Sema3E-plexinD1 signalling in CD11c+ DC is a critical cascade that modulates AHR, airway inflammation, and tissue remodelling in the HDM model of asthma.

## Abbreviation

AHR: airway hyperresponsiveness
AM: alveolar macrophage
ASM: airway smooth muscle cell
a-SMA: a-smooth muscle actin
BALF: bronchoalveolar lavage fluid
BM: bone marrow
BMDM: bone marrow-derived macrophage
DC: dendritic cell
GATA3: GATA binding protein 3
HDM: house dust mite
IM: interstitial macrophage
IRF-4: interferon regulatory factor 4
KO: knockout
mLN: mediastinal lymph node
PDL2: programmed death-ligand 2
qRT-PCR: quantitative real-time PCR
RORγt: Retinoic Acid Receptor-Related Orphan Receptor Gamma T
Sema3E: semaphorin3E
Treg: regulatory T cell
WT: wild-type.

## Acknowledgment

The authors would like to thank Dr. Christine Zhang (Flow Cytometry Core Facility, University of Manitoba) for her help on flow cytometry experiments.

